# Developmental underpinnings of morphological disparity in the avian bony palate

**DOI:** 10.1101/2025.07.22.666138

**Authors:** Olivia Plateau, Guillermo Navalón, Juan Benito, Daniel J. Field

## Abstract

The deepest phylogenetic divergence in crown birds gave rise to Palaeognathae and Neognathae, clades exhibiting divergent bony palate morphologies. This observation led to the longstanding hypothesis that the distinctive palate arrangement of palaeognaths retains the ancestral crown bird condition, but recent fossil evidence instead suggests that aspects of palaeognath palate morphology are derived from a neognath-like ancestral condition. While neoteny has been hypothesised as a mechanism underpinning the distinctive palatal morphologies of palaeognaths, this hypothesis has never been tested with a broad phylogenetic assessment of morphological variation through avian palate ontogeny. Here, we quantitatively assess morphological variation of the palate through the post-hatching ontogenies of 70 bird species representing all major extant phylogenetic subclades, clarifying the ontogenetic mechanisms giving rise to avian palate disparity. Though palaeognaths exhibit distinct ontogenetic changes in the PPC relative to neognaths, we find no signatures of heterochrony—including neoteny—underlying these developmental differences. However, important patterns of morphological change in the avian palate appear to be dictated by variation in developmental mode. Our results document the effects of post-hatching development on a key morphofunctional system in the avian skull, and, more broadly, highlight the influence of developmental mode on morphological evolvability across crown group birds.

## Introduction

The deepest phylogenetic divergence within crown birds, between palaeognaths (ratites and tinamous) and neognaths (all other birds), was originally recognised on the basis of the morphology of the skull’s pterygoid-palatinum complex (hereafter, PPC)^1–4^. The PPC is composed of five bones: an unpaired vomer and paired palatines and pterygoids. Whereas palaeognaths exhibit a fused, immobile contact between the pterygoid and palatine, the PPC of neognaths is characterised by a mobile joint between these elements, enabling an enhanced degree of palatal kinesis^1,4,5^.

Multiple aspects of palaeognath morphology, including their distinctive PPCs, have been hypothesised to reflect the retention of the ancestral crown bird condition^6–9^. By contrast, certain authors have suggested that the apparently plesiomorphic aspects of the palaeognath PPC may in fact be derived, reflecting a hypothesised paedomorphic developmental shift from a neognath-like ancestral condition associated with the loss of flight^10–12^. Towards reconciling these alternative views, the PPC morphologies of Mesozoic avialans (i.e., *Archaeopteryx, Gobipteryx* and *Hesperornis*) have been investigated to illuminate the nature of the PPC along the crownward portion of the avian stem lineage. Early investigations^13^ concluded that the PPCs of Palaeognathae share more similarities with those of Mesozoic stem birds than with Neognathae, bolstering the hypothesis that palaeognaths retain a more plesiomorphic arrangement of the palate than neognaths. However, unanticipated paleontological discoveries have recently added additional nuance to this debate, with new specimens of Mesozoic near-crown stem birds recognised as exhibiting mobile PPC arrangements similar to those of some neognaths ^14–17^.

Despite the potential importance of these palaeontological findings, to date they lack formal corroboration from developmental datasets beyond investigations limited to a narrow phylogenetic sample ^18,19^. Here, we investigate post-hatching ontogeny of the PPC complex across a broad range of extant birds (family-level representation across Palaeognathae and members of all major neognath subclades; Figs. 1, 2) to clarify the developmental and evolutionary underpinnings of morphological variation in the avian PPC. As the vomer is often reduced and vestigial in neognaths ^20–24^, we focus on morphological variation of the palatine and the pterygoid. We use landmark-based three-dimensional geometric morphometrics to quantify morphological changes in the PPC throughout ontogeny and assess morphological variation within and among major clades to assess the main ontogenetic drivers of morphological variation between Palaeognathae and Neognathae. These data provide unprecedented insight into the nature of morphological variation in the avian PPC and clarify patterns of evolutionary change in the PPC near the origin of crown birds.

**Figure 1.**
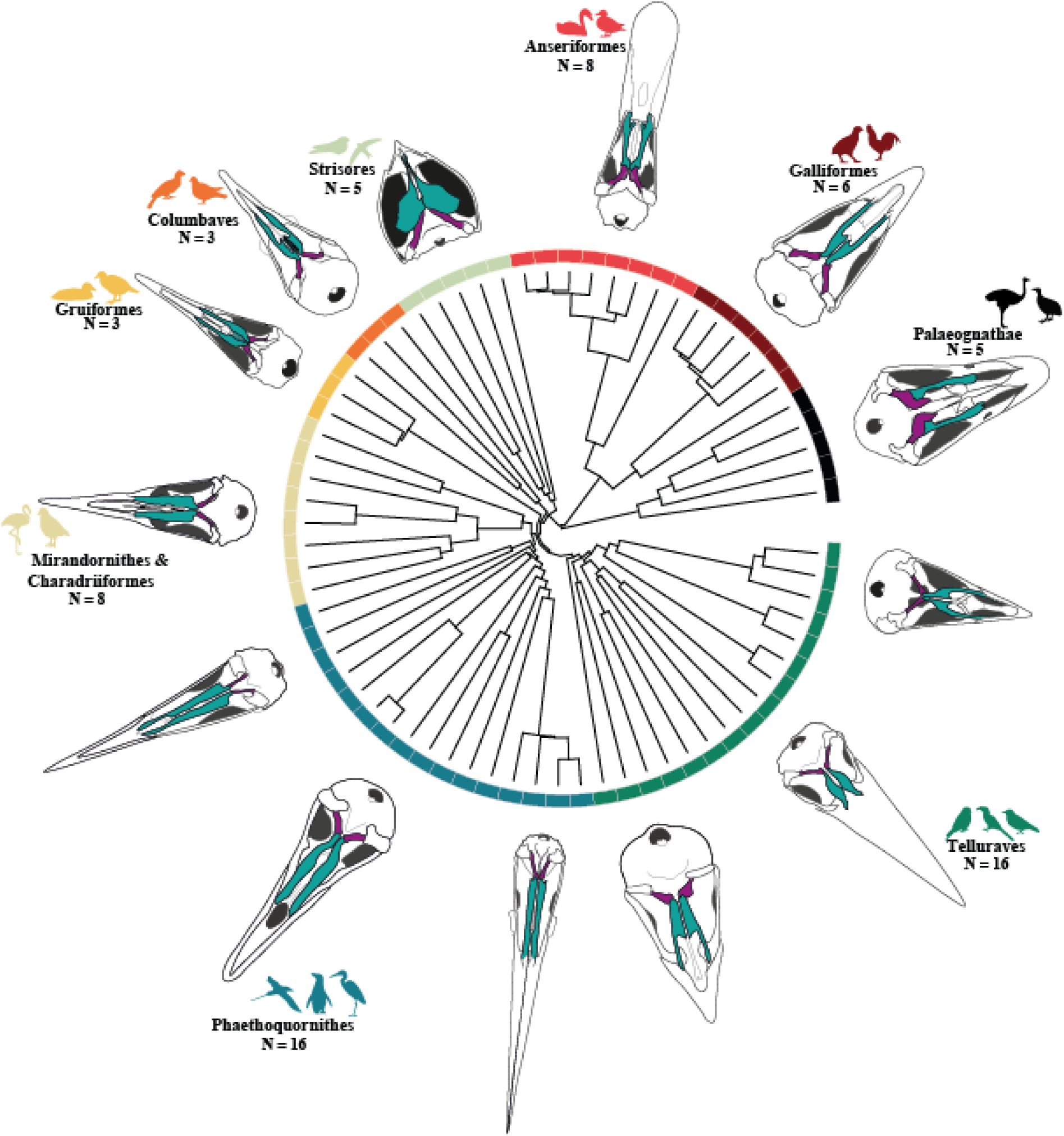
The pterygoid-palatinum complex (PPC) across bird phylogeny. Colours on the phylogeny represent the nine subgroups compared in our investigation, following the topology of Prum et al. (2015). Reconstructions of the skull in ventral view are illustrated for each subgroup under investigation, highlighting the bones of the PPC (palatine in teal, pterygoid in purple). From Palaeognathae to Telluraves, counterclockwise, the species illustrated are *Struthio camelus* (Palaeognathae), *Gallus gallus* (Galliformes), *Anas platyrhynchos* (Anseriformes); *Nyctibius griseus* (Strisores); *Columba livia* (Columbaves); *Gallirallus australis* (Gruiformes); *Burhinus senegalensis* (Charadriiformes); *Gavia stellata* (Phaethoquornithes); *Macronectes halli* (Phaethoquornithes); *Ardea cinerea* (Phaethoquornithes); *Opisthocomus hoazin* (sister to Telluraves); *Pteroglossus viridis* (Telluraves); and *Pica pica* (Telluraves). See Supplementary Data 9for detailed information on specimens. Reconstructions not to scale.

**Figure 2.**
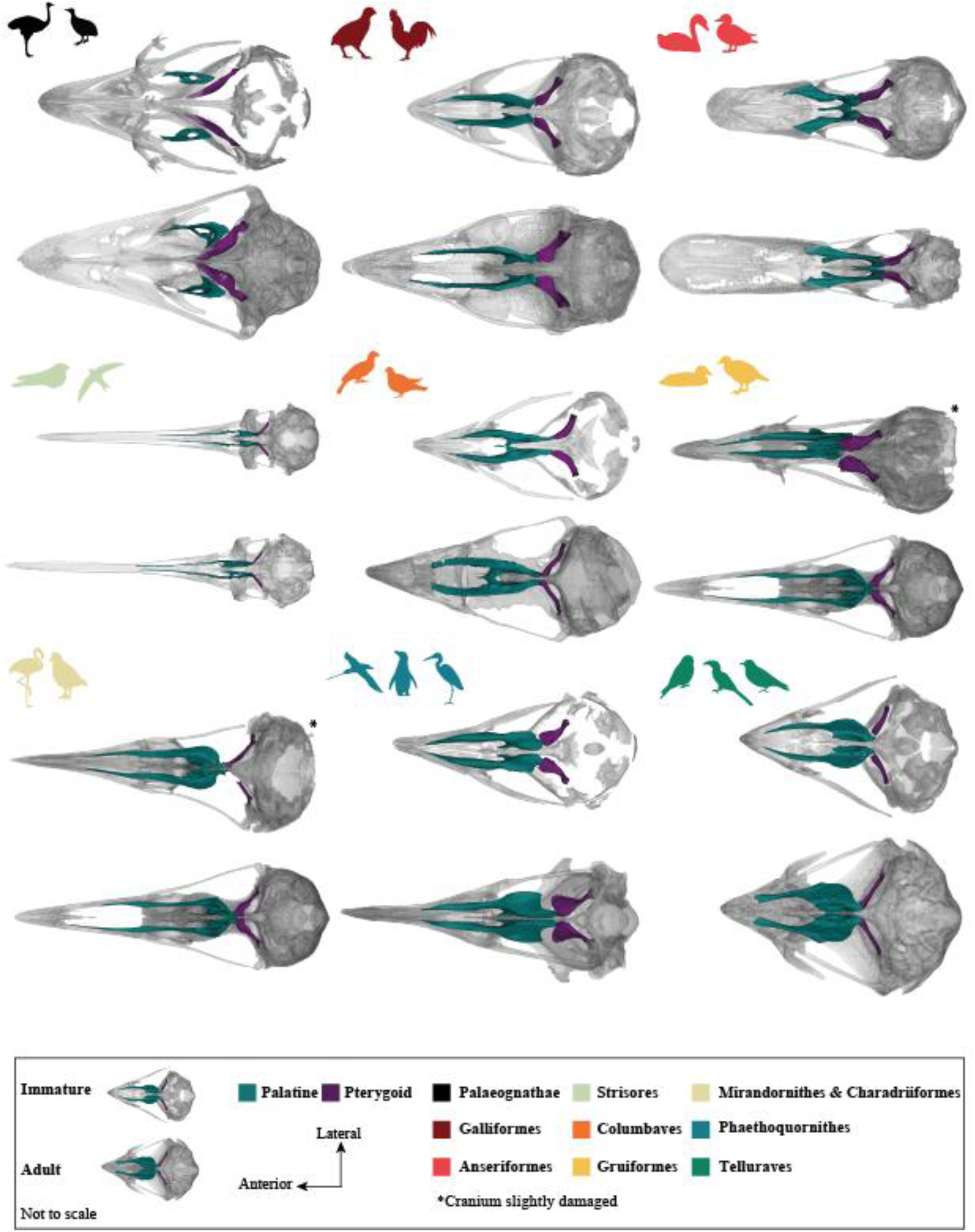
Ontogenetic comparative morphology of the palatine and pterygoid of the primary extant bird clades investigated. For each group we illustrate the cranium of an immature and an adult specimen in ventral view. Species illustrated are: *Dromaius novaehollandiae* (Palaeognathae); *Gallus gallus* (Galliformes); *Anas platyrhynchos* (Anseriformes); *Patagona gigas* (Strisores); *Tauraco erythrolophus* (Columbaves); *Fulica americana* (immature) / *F. atra* (adult; Gruiformes); *Fratercula arctica* (Charadriiformes); *Spheniscus demersus* (Phaethoquornithes); *Falco naumanni* (Telluraves). See Supplementary Data 9 for full information about specimens investigated.

## Results

### Quantifying morphological disparity among major bird clades

We compared macroevolutionary and ontogenetic variation in PPC morphology among major clades of extant birds in two ways: 1) treating the PPC as a single complex, and 2) examining shape variation of the pterygoid and palatine as individual elements.

When considering the PPC as a single complex, the morphospace defined by the first two PCs accounts for 52% of PPC shape variation (Fig. 3). The negative end of PC1 is associated with anteroposteriorly short but mediolaterally wide palatines combined with antero-posteriorly elongate pterygoids with a mediolaterally broad and robust quadrate articulation, whereas the positive end of PC1 captures anteroposteriorly elongate palatines with anteroposteriorly short pterygoids (Fig.3A). The negative end of PC2 is associated with anteroposteriorly elongate palatines and pterygoids, whereas the positive end of PC2 captures anteroposteriorly short but mediolaterally wide palatines combined with anteroposteriorly short pterygoids (Fig. 3A).

**Figure 3.**
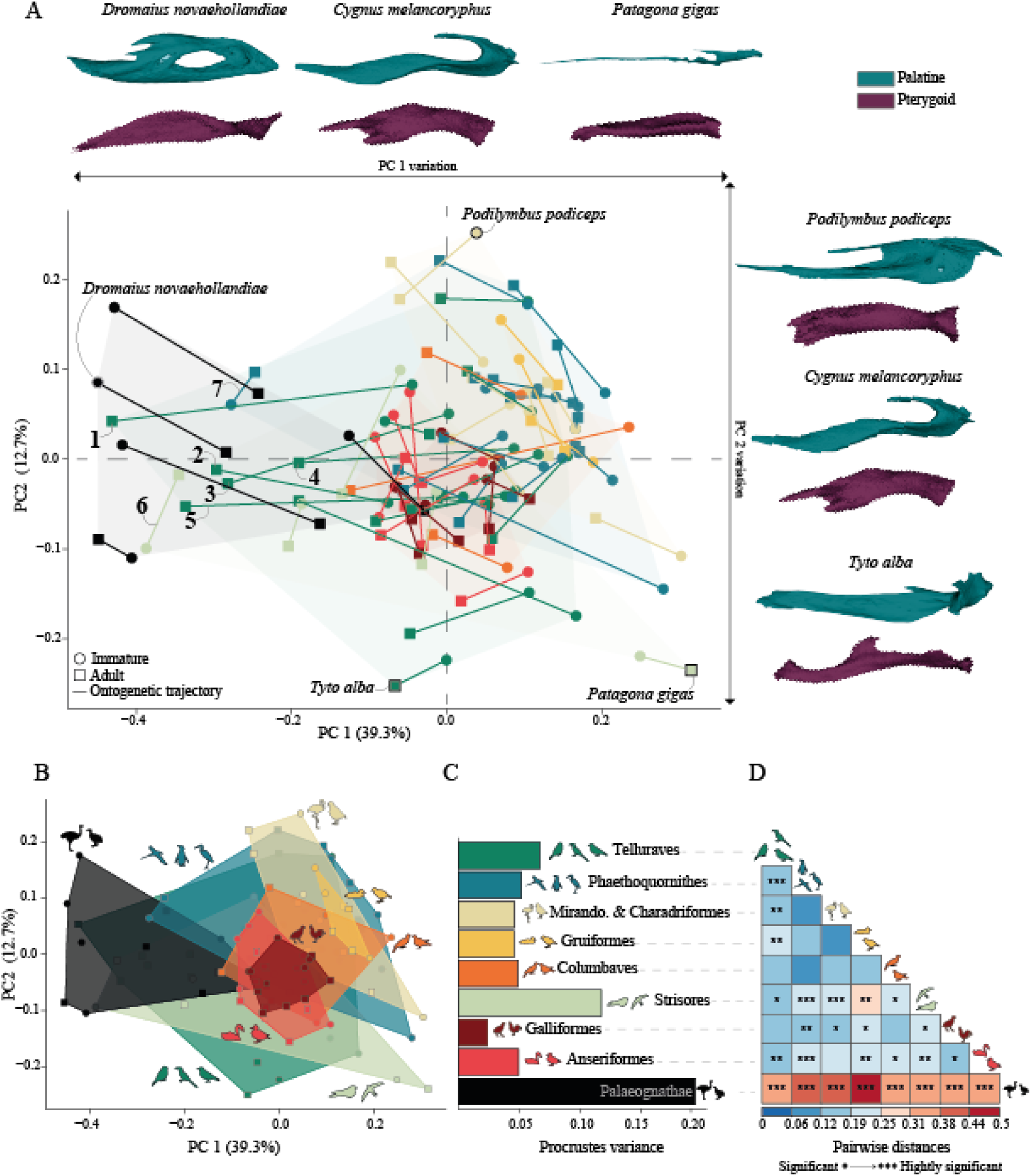
Principal Component Analysis (PCA), morphological disparity and pairwise group comparisons of pterygoid-palatinum complex (PPC) ontogenetic disparity across crown birds. A) PPC morphospace based on the first two principal components. B) Bar plot showing Procrustes variance per major subclade. C) Correlation plot of pairwise distances between each subclade, with stars indicating the levels of significance in each comparison (* = 0.05>P>0.01; ** =0.01>P>0.001; *** 0.001>P). For A, B and C subclade colours follow Figure 1. In A, lines link immature and adult specimens of the same species; specific numbers in the PCA correspond to 1: *Ara ambiguus*; 2: *Lorius garrulus*; 3: *Cacatua alba*; 4: *Pteroglossus viridis*; 5: *Anorrhinus galeritus*; 6: *Podargus strigoides*; 7: *Phaethon lepturus*. See Supplementary Data 9 for detailed information on specimens. Reconstructions not to scale (A).

Palaeognathae and Neognathae are differentiated along PC1 (Fig. 3A-B), with palaeognaths exhibiting considerably greater morphological disparity than neognaths (Fig. 3C), including several of the most divergent PPC morphologies of any extant birds (Fig. 3D; Supplementary Data 1 and 2). While the palatines of some palaeognaths (i.e. ostrich and the tinamou *Crypturellus*) are anteroposteriorly elongate and mediolaterally narrow, others (i.e. emu, rhea, and the tinamou *Nothoprocta*) are much shorter and wider. In contrast, whereas the pterygoids of emu, ostrich and rhea are anteroposteriorly short and mediolaterally broad, those of tinamous are elongate and tube-shaped.

Within Neognathae, the subclades Strisores (nightbirds) and Telluraves (landbirds) exhibit the greatest range of morphological disparity, and overlap with palaeognaths along PC1 and PC2 (Fig. 3B-C). However, these instances of overlap (Fig. 3A) are restricted to a small handful of neognaths (7 species out of 70) with especially derived cranial and PPC morphologies (adult parrots, toucans, hornbills and frogmouths, and both immature and adult tropicbirds), all of which exhibit relatively short palatines and extremely wide pterygoids. Certain divergent palaeognath morphologies also result in overlap with neognaths along PC1 and PC2; for instance, the adult ostrich overlaps with Galloanserae (land- and waterfowl; Fig. 3A-B).

Nonetheless, all neognath subclades exhibit substantial overlap along PC1-PC2 (see SI, Supplementary Results). Although pairwise distances are relatively low among neognath subclades, statistically significant morphological differences distinguish Galloanserae, Strisores, Phaethoquornithes and Telluraves from other subclades (Fig. 3C-D; Supplementary Data 1 and 2).

Analysing the palatine and pterygoid as individual elements (Fig. 4) yields similar general patterns along PC1and PC2 to those from our analyses of the PPC as a single complex.

**Figure 4.**
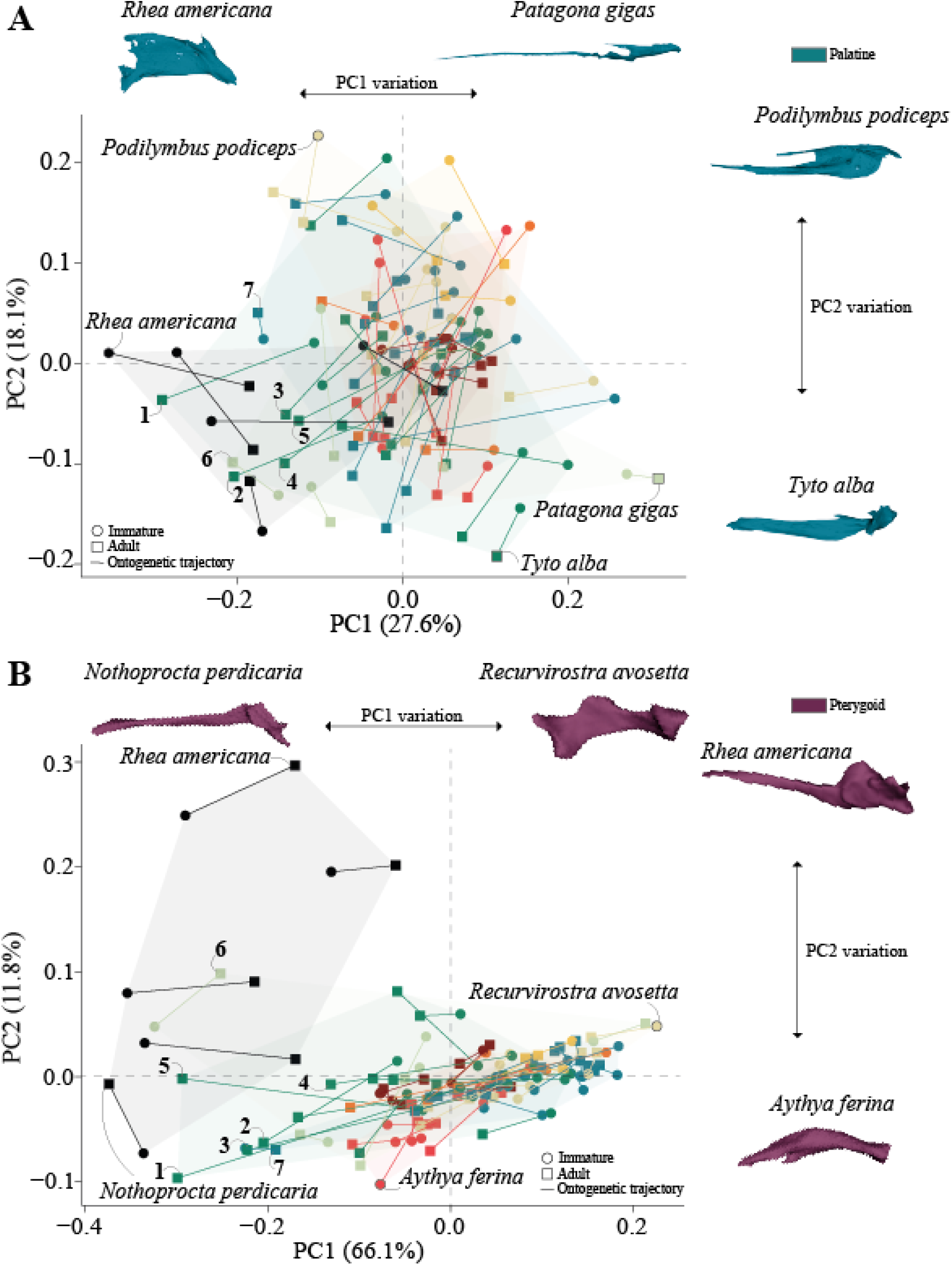
Principal Component Analysis (PCA) of palatine and pterygoid morphology. A) Palatine morphospace based on the first two PC scores. B) Pterygoid morphospace based on the first two PC scores. Subclade coloursfollow Figure 1. Circles indicate immature specimens and squares indicate adult specimens; each line links immatures and adults of the same species. Specific numbers in the PCA correspond to 1: *Ara ambiguus*; 2: *Lorius garrulus*; 3: *Cacatua alba*; 4: *Pteroglossus viridis*; 5: *Anorrhinus galeritus*; 6: *Podargus strigoides*; 7: *Phaethon lepturus*. See Supplementary Data 9 for detailed information on specimens. Reconstructions not to scale.

Collectively, the first two principal components account for 46% of total shape variation for the palatine (Fig. 4A), and 78% for the pterygoid (Fig. 4B). Furthermore, geometric dissimilarity between the pterygoids of Palaeognathae and Neognathae is greater than that for the palatines (Extended Data Fig. 1). In contrast, within Neognathae, palatine shape is more distinct among major clades than the pterygoid, especially among Anseriformes, Mirandornithes and Charadriiformes, and between Telluraves and Strisores (Extended Data Fig. 1).

Dividing our total sample into immature and adult subsets reveals patterns of morphological disparity that are broadly similar to those of the combined dataset (Extended Data Fig. 2,3; Supplementary Data 3). However, both in terms of within-clade morphological disparity and among-clade pairwise distances, immature palaeognaths are more morphologically variable than adults are. In contrast, most neognath adults are more disparate than immatures are (Extended Data Fig. 2,3; Supplementary Data 3). Nevertheless, differences among clades in the whole dataset are greater than those in the separate immature and adult subsets. The immature and adult subsamples exhibit comparable phylogenetic signal, with only slight variation in *Kmult* (Immature: K = 0.78, P = 0.001; Adult: K = 0.83, P = 0.001).

### Ontogenetic trajectories across avian phylogeny

To compare ontogenetic variation within and among major extant bird clades (Fig. 1), we calculated angles between the ontogenetic shape trajectories of each species and estimated the degree of ontogenetic divergence (i.e. individuals becoming less similar throughout ontogeny) *versus* convergence (i.e. individuals becoming more similar throughout ontogeny) of each pair of species based on pairwise Procrustes distances between immature and adult individuals (see Methods). To identify heterochronic shifts in ontogenetic variation among groups (i.e. changes in the rate and timing of developmental processes^25^), hypothesised as a driver of the distinct palate morphologies of Palaeognathae and Neognathae^26^, we applied a multi-test approach comparing size-shape relationships among major clades (see Methods).

### Group-specific shape differences in ontogenetic trajectories

Within Palaeognathae, comparisons of the angle of ontogenetic trajectories show a bimodal distribution (Fig. 5A), with a cluster of taxa around 60° (the tinamou *Nothoprocta* vs. all other palaeognaths), and a second cluster around 100° (comparisons among all other palaeognaths). Neognaths exhibit a greater range of differences in ontogenetic trajectories, (31° - 137°), with a continuous distribution of values between the extremes represented by the trajectories of *Recurvirostra avosetta* vs *Plegadis falcinellus* and *Falco tinnunculus* vs *Spheniscus demersus*. (Fig. 5A). When Palaeognathae and Neognathae are compared, angles among ontogenetic trajectories exhibit a continuous distribution from 60° (*Perdix perdix* vs *Dromaius novaehollandiae*) to 137° (*Crypturellus tataupa* compared vs *Plegadis falcinellus*) (Fig. 5A).

**Figure 5.**
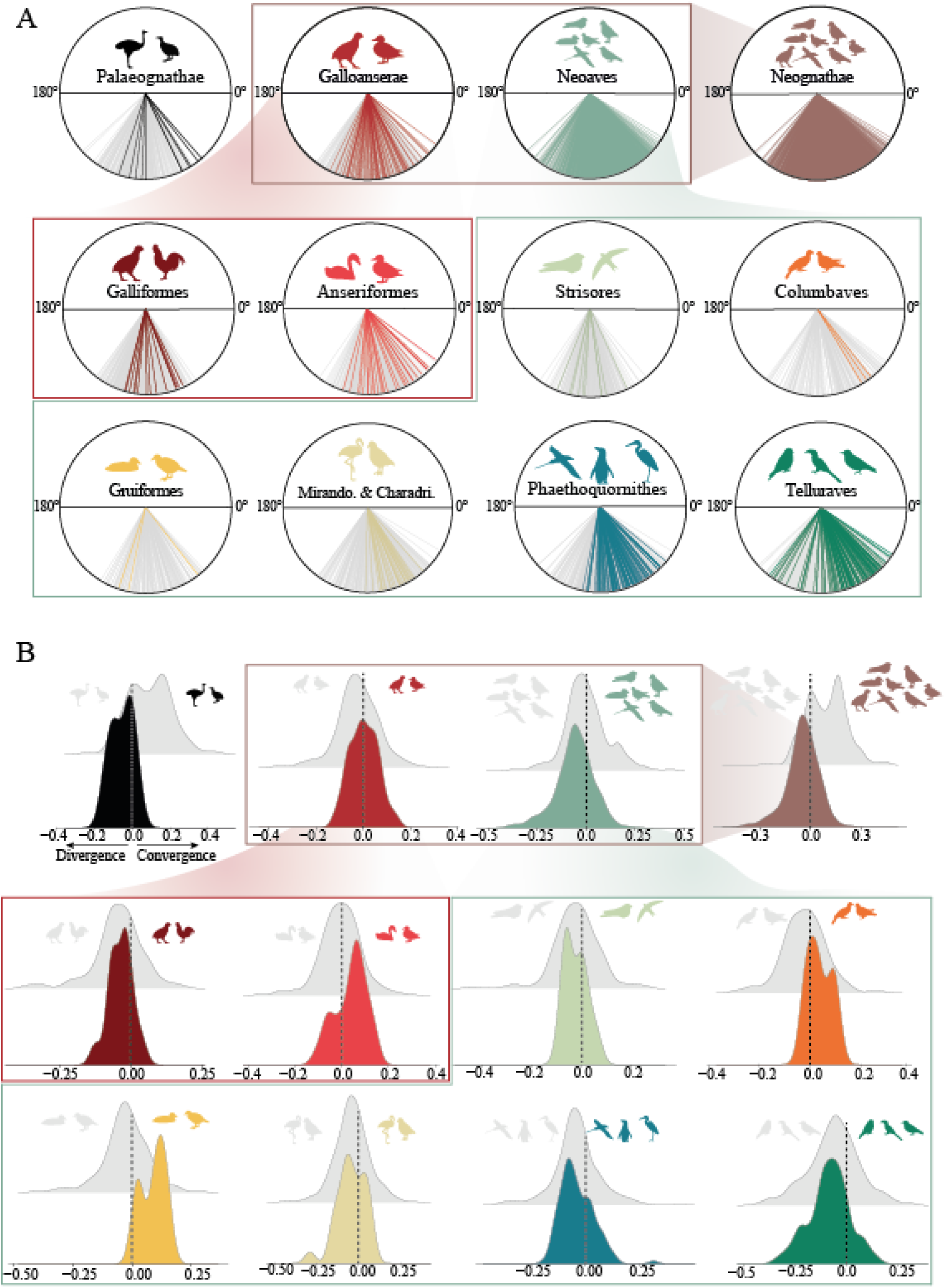
Comparisons of ontogenetic trajectories between and within major clades of crown birds. A) Circular plot of inter-species angles within and between groups. B) Frequency plot of inter-species divergence/convergence values within and between groups. For A and B subclade colours follow Figure 1 for Palaeognathae, Galliformes, Anseriformes, Strisores, Columbaves, Gruiformes, Mirandornithes & Charadriiformes, Phaethoquornithes, and Telluraves.

Within the major subclades comprising Neognathae, comparisons within Galloanserae exhibit a narrower range of variation (between 37°, when comparing *Dendrocygna arborea* to *Malacorhynchus membranaceus*, to 117°, when comparing *Anas platyrhynchos* to *Numida meleagris*), and a more heterogeneous distribution of values than comparisons within Neoaves (Fig. 5A). The greater disparity observed in Neoaves is largely attributable to variation within the hyperdiverse subclade Telluraves, which range from 33° (*Anorrhinus galeritus* vs *Pteroglossus viridis*) to 127° (*Falco tinnunculus* vs *Falco naumanni*). Other neognath subclades occupy essentially the full range of variation between the extremes delimited by Palaeognathae and Telluraves (see SI, Supplementary Results for more details). Between clades, however, a broad spectrum of variation in ontogenetic trajectories is evident, with almost uniform variation between 31° and 137° (Fig. 5A). Except for especially high values of intraclade variation in Telluraves, the most extreme differences are restricted to interclade comparisons rather that arising from intraclade comparisons.

### Ontogenetic shape divergence and convergence among birds

Within most major extant bird subclades, we find a general pattern in which the PPC morphologies of different species diverge throughout ontogeny (Fig. 5B); that is, immatures of different species exhibit comparatively similar PPC morphologies whereas adults are more geometrically distinct (although see SI, Supplementary Results for more details). This pattern is further exaggerated in Telluraves, which exhibit the greatest degree of within-clade ontogenetic shape divergence (Fig. 5B; Supplementary Data 5). Notably, Anseriformes exhibit the opposite pattern, with interspecific PPC morphology converging throughout ontogeny (Fig. 4C), while Galliformes show more ontogenetic divergence than Anseriformes (see SI, Supplementary Results for more details).

Among major neognath subclades, PPC shape tends to diverge throughout ontogeny. By contrast, palaeognath PPC morphology converges upon adult neognath PPC morphology as ontogeny progresses (Fig.5B, Extended Data Fig.4), in accordance with our finding that differences in the shape of the PPC between palaeognaths and neognaths are greatest at early ontogenetic stages (Extended Data Fig. 2). These morphological differences are more pronounced in the pterygoid than in the palatine (Extended Data Fig. 2).

### Non-heterochronic ontogenetic variation in the PPCs of Palaeognathae and Neognathae

Based on model comparisons (see Materials and Methods), we found that comparisons of ontogenetic trajectories between Palaeognathae and Neognathae were the most effective basis for testing heterochronic hypotheses for the full PPC complex as well as for the palatine in isolation (Extended Data Fig.4A, B). However, for the pterygoid, the best model for testing for heterochrony instead involved comparing ontogenetic allometries between Palaeognathae and the major subgroups of Neognathae (Extended Data Fig.4C).

For both the isolated palatine and the PPC complex, palaeognath allometric ontogenetic slopes differ significantly from Neognathae (Extended Data Fig.4A, B, Supplementary Data 6). Non-heterochronic processes appear to underly ontogenetic changes between palaeognaths and neognaths (Supplementary Data 6, Thf2: significant), and their trajectories tend to converge^27,28^ as post-hatching ontogeny progresses (Extended Data Fig.4A, B) as adults show fewer morphological differences than immatures. For the pterygoid, excluding the comparison between Palaeognathae and Galliformes (see below), allometric ontogenetic slopes of Palaeognathae also significantly differ from those of Neognathae (Extended Data Fig.4C, Supplementary Data 6). Furthermore, these trajectories also differ in shape trajectory (Supplementary Data 6, Thf2: significant)) and their ontogenetic shape trajectories tend to converge towards each other during post-hatching ontogeny (Extended Data Fig.4C). While ontogenetic allometric trajectories between Palaeognathae and Galliformes show no differences in slope, their intercepts are significantly different (Supplementary Data 6, Thf1: significant), indicating parallel slopes, rather than heterochronic variations from a common ancestral ontogenetic allometric trend. Overall, our results illustrate a general lack of evidence for heterochronic changes giving rise to the morphological differences in the PPCs of Palaeognathae and Neognathae (see SI, Supplementary Results for more details).

### Influence of developmental mode on ontogenetic variation in the avian PPC

Finally, we interrogated how variation in PPC ontogenetic trajectories is shaped by variation in life history strategies along the altricial-precocial spectrum^29^(Fig. 6A). This provides an additional exploration of the macroevolutionary effects of developmental variation beyond linear relationships between size and shape across ontogenies. Across extant birds we found a continuous range of variation describing ontogenetic divergence and convergence and note that altricial taxa exhibit markedly greater degrees of ontogenetic divergence than precocial taxa (Fig. 6A; Supplementary Data7). To statistically assess the relationship between life history traits and ontogenetic variation (changes in PPC shape and morphospace trajectory), we assessed the correlation between our ontogenetic parameters (morphological distance, angles between ontogenetic trajectories, and *Divergence-Convergence* values) and a semi-continuous quantitative index of avian developmental mode^30^. The resultant Ordinary Least Squares linear models (Fig. 6 B-C; Supplementary Data 8) revealed weak but significant correlations between all variables (i.e., Distance: *R*^2^ = 0.11, P = *0.024*; Angle: *R*^2^ = 0.08, P = *0.044*; *Divergence – Convergence*: *R*^2^ = 0.14, P = *0.07*) that became non-significant when phylogenetic relationships were controlled for (Fig. 6B). This pattern (i.e., OLS significant, PGLS non-significant) may indicate that shifts in PPC ontogenetic variables and developmental mode both occurred early in the evolutionary history of crown birds (Clauss et al., 2013, Fig. 6), at the base of large clades and not within these subclades (Fig. 6).

**Figure 6.**
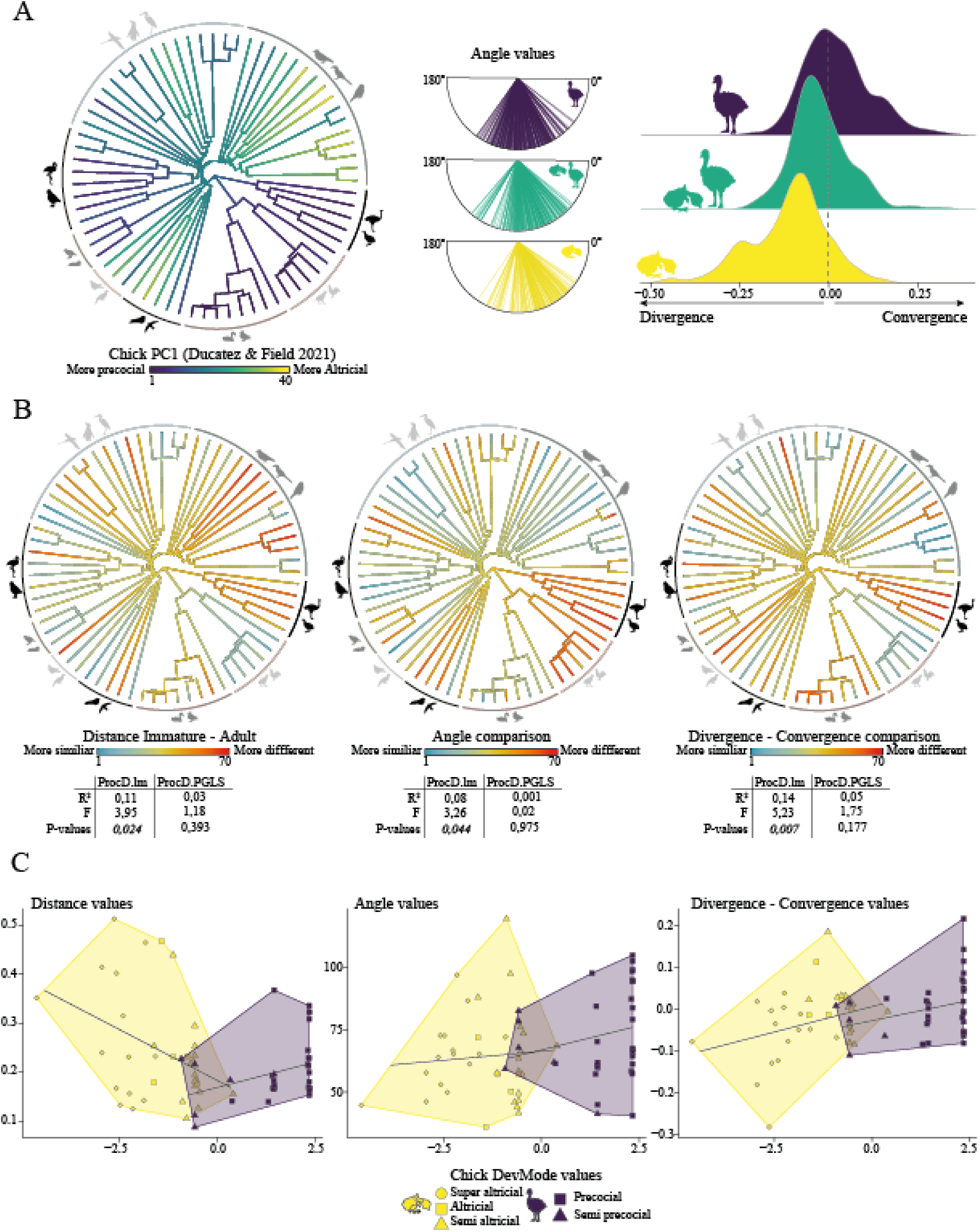
Avian developmental mode and its imprint on ontogenetic variation in the avian bony palate. A) Plot of Chick PC1 (Ducatez and Field (2021)^26^) across avian phylogeny (left). Chick PC1 corresponds to the first axis of the PCA performed on a matrix of seven discrete developmental traits (hatchling down coverage, hatchling eye condition, age at eye opening, hatchling locomotor activity, hatchling feeding capacity, ratio of time spent in the nest to age at first flight, and post-nest behavior), capturing the atricial-precocial spectrum (Ducatez and Field (2021)^26^. Circular plot of inter-species angles (centre) and frequency plot of inter-species divergence/convergence relative to developmental mode categories (right). B) Distance map between immature and adult (left), angle (middle) and divergence – convergence (right) values between individual species and estimated ancestral state as a reference value (see Materials and Methods) across avian phylogeny. Circular lines in different shades of gray represent the nine avian subclades under investigation; see Figure 1. Tables under each phylogenetic tree indicate statistical results of OLS and PGLS regressions between ontogenetic parameters and developmental mode (Chick PC1). C) Plot of the OLS regression (black line) between immature -adult distances (left), angles (centre), and Divergence-Convergence (right) values and Chick PC1 values.

## Discussion

Our study represents the most detailed quantitative exploration of avian PPC ontogeny to date and fills an important gap in our understanding of avian palate evolution.

We found no evidence for signatures of heterochronic processes (that is, evolutionary changes in the rate and timing of developmental changes) contributing to the differentiation of diagnostic palate morphologies of Palaeognathae and Neognathae, in agreement with more limited previous explorations^18^. However, we found evidence that variation in developmental mode plays a prominent role in influencing PPC ontogeny across avian diversity. Specifically, precocial birds tend to exhibit convergent morphological trajectories of the PPC throughout ontogeny, whereas the PPCs of altricial groups tend to diverge as ontogeny progresses (Fig.6A). These results mirror patterns from other anatomical modules across the avian body^31^, and suggest that developmental mode variation may modulate the strength of constraints on PPC morphology at macroevolutionary scales.

Moreover, our results suggest that major shifts in developmental mode and correlated large-scale patterns in PPC ontogenetic trajectories occurred only a limited number of times, early in the evolutionary histories of major crown bird clades, complicating straightforward inferences regarding PPC morphology in the ancestral crown bird from extant taxa. Our results hint at an evolutionary scenario in which rapid evolutionary changes in developmental mode in the early Cenozoic were associated with major evolutionary changes in PPC morphology, overprinting plesiomorphic morphological signatures.

Clear morphological differences in the PPC between Palaeognathae and Neognathae are discernible from analyses of both individual PPC components and the entire complex. In all versions of our analyses, Palaeognathae shows greater morphological disparity than all the major clades within Neognathae. Individually, the pterygoid shows a greater degree of morphological differentiation between these two major clades than the palatine, reflecting a notable degree of constraint on the pterygoid morphology of neognaths and a lack thereof in palaeognaths.

These interclade patterns reflect fundamental differences in the configuration and function of the avian palate. We hypothesise that the fusion and loss of mobility between the palatine and the pterygoid in Palaeognathae^2,32^ may have released ancestral functional constraints associated with cranial kinesis, allowing these bones to vary more freely. This effect may have been less pronounced in the palatine than in the pterygoid due to the palatine’s immobile connections with the rostrally positioned premaxilla and the laterally placed maxilla (and, when present, the medially positioned vomer ^21–23^). By contrast, in all prokinetic birds the pterygoid articulates only at mobile joints (with the palatine rostrally and the quadrate caudally)^2,4,22,33,34^.

Within Neognathae, the morphology of the pterygoid is remarkably conservative, presumably related to the maintenance of its key prokinetic role within this clade as a force-transmitting lever between the quadrate and palatine. By contrast, the palatine is considerably more variable, with Strisores (nightbirds) and Telluraves (landbirds) exhibiting the greatest levels of morphological disparity in the palatine, in line with these clades exhibiting some of the most extreme rostral morphologies among extant birds^35–37^. Given that the major components of the rostrum appear to ossify at relatively similar stages of development^19,38–41^, we suggest that the palatine may be more affected by adaptive changes in rostral morphology than the pterygoid, which warrants future investigation in studies of avian pre- and post-hatching cranial development.

This hypothesis may also be supported by myological evidence. Previous investigations have highlighted the impact of muscle arrangement on the skull shape of other vertebrates, such as mammals^42,43^. Although these relationships are considerably less well-studied in birds, the *musculus pterygoideus* (*Dorsalis* and *Ventralis*) and the *musculus protactor pterygoidei et quadrati*^33^ are associated with avian beak opening and closing, connecting the palatine to the pterygoid, and the pterygoid to the quadrate, respectively^2^. The arrangement of these muscles is relatively simple in Palaeognathae^33,44,45^ and more complex in Neognathae, with numerous muscle subdivisions increasing the number of muscle-bone contacts^46–48,33^. The more complex muscular attachments associated with the neognath pterygoid may be related to its comparatively constrained degree of morphological disparity relative to that of palaeognaths, and relative to the neognath palatine.

Notably, some neognaths overlap with palaeognaths along the main axes describing PPC geometry, such as some parrots (Psittaciformes), frogmouths (Strisores: Podargidae), hornbills (Bucerotiformes), and toucans (Piciformes: Ramphastidae). This pattern appears to be driven primarily by palatine morphology rather than the pterygoid (Fig.3A, Fig.4A-B), and these taxa generally exhibit short but extremely wide palatines, a condition widespread in Palaeognathae. Importantly, these taxa include an array of neognaths with secondarily reduced palatal kinesis (e.g., toucans^49^), as well as kinetic taxa with divergent rostral morphologies (e.g., parrots^50^), suggesting that multiple factors may be responsible for these convergent shifts in different clades.

We note that limitations of current morphometric tools represent a major challenge to quantifying what is perhaps the most fundamental cranial distinction between palaeognaths and neognaths: the fused connection between the pterygoid and palatine of palaeognaths^1–3,32^. For instance, the general arrangement of the palatine and pterygoid in the frogmouth *Podargus strigoides* is strikingly similar to that of palaeognaths (note overlapping PPC geometries along PC1 and PC2 in Fig. 3A and Fig. 4B). Although the anterior articular surface of the pterygoid sits atop the dorsal surface of the caudal palatine in *Podargus*, these elements remain unfused, unlike the condition in Palaeognathae where these elements fuse to one-another.

Clade-specific differences in palate morphology are already apparent at the hatchling stage in most bird lineages. For instance, differences in form between palaeognath and neognath PPCs and pterygoids are even more pronounced among hatchlings than adults. Whereas palaeognaths exhibit their greatest degree of morphological disparity in the PPC at the hatchling stage, neognaths (with the exception of Anseriformes) exhibit greater disparity at the adult stage than at the hatchling stage (see SI, Supplementary Discussion for a more detailed discussion). These patterns appear to be dictated in part by variation in developmental mode. Specifically, in precocial and super precocial^29,51^ taxa (e.g., all palaeognaths, anseriforms) a great deal of ossification and morphological change occurs during pre-hatchling development. By contrast, altricial and super altricial^29,51^ taxa (all of which are neognaths) generally hatch in a comparatively weakly ossified state, in which interactions between muscle action and extensively unossified skeletal elements can induce pronounced variation in bone morphology during post-hatching growth^31^. Galliformes, the precocial and super precocial sister taxon to Anseriformes, represent a notable exception to this pattern, exhibiting the lowest observed values of morphological disparity at both stages. This observation is in line with the generally conservative cranial morphology of galliforms^52^, hinting at additional clade-specific developmental constraints^41^ responsible for canalising galliform cranial morphology that are worthy of further investigation. Crucially, only Neoaves experience post-hatching palatal segmentation (see Online Methods), which is absent in Palaeognathae and incomplete in Galleoanserae^21,32,53,54^, yet the role played by this ontogenetic transformation in shaping palatal morphology is uncertain.

Whereas altricial taxa tend to exhibit divergent ontogenetic trajectories of PPC morphology, and precocial taxa exhibit convergent trajectories (Fig.6A), semi-altricial and semi-precocial taxa tend to exhibit intermediate values that are challenging to fully disentangle. This recalls broader macroevolutionary patterns among birds and other vertebrates, in which clear ecomorphological patterns may only be discernible at the extremes of ecological variation, in which stronger selective pressures yield consistent morphological solutions. For instance, beak shape—a well-studied aspect of avian skull variation—is distinct among taxa exhibiting highly distinct ecologies, but otherwise exhibits extensive overlap in form^55,56^.

Previous studies have reported evidence for the contribution of heterochronic processes to craniofacial evolution near the origin of modern birds^57–59^ and within major lineages such as

Strisores^36^. However, our results suggest limited evidence for heterochrony as a major driver of morphological divergence in the avian PPC, emphasising that inferred macroevolutionary patterns in ontogenetic allometry may differ not only among anatomical elements but also in line with the phylogenetic scale under investigation ^60–62^.

Though long challenging to investigate quantitatively^30^, previous research has illustrated the influence of the precocial-altricial spectrum on patterns of variation across the avian crown group, influencing biological parameters as varied as genomic evolutionary mode^63^ and anatomical modules such as the legs^31^, brain^64^, and beak^55,56^. Collectively, these studies and our present results suggest that the evolutionary emergence of avian altricial development was associated with an increase in morphological evolvability, perhaps representing a key innovation contributing to the striking imbalance of diversity and disparity of form that distinguishes precocial and altricial bird taxa in the present day. The underappreciated influence of developmental mode variation on macroevolutionary shifts in skeletal form highlights the need for further research into the evolutionary origins of avian altriciality and its phenotypic consequences.

## Online Methods

### Sampling, and 3D Data acquisition

To quantify morphological variation in the Pterygoid-Palatinium Complex of extant birds, we sampled species of all major bird lineages and grouped them following major clades in the Prum et al., 2015 phylogeny [Palaeognathae (n=5), Galliformes (n= 6), Anseriformes (n= 8), Strisores (n= 5), Columbaves (n= 3), Gruiformes (n= 3), Mirandornithes and Charadriiformes (n= 8), Phaethoquornithes (n= 16) and Telluraves + Hoatzin (Inopinaves, n= 16). One immature and one adult were sampled for a total of 70 extant bird species (Supplementary Data 9). Although data on absolute ontogenetic age is often missing from museum species, we sampled specimens as close as possible to the hatching stage based on size, plumage characteristics and the state of skull bone fusion^24^. For some Palaeognathae, specimens treated as adults in our analyses were necessarily represented by subadults whose skull shape and size are almost identical to those of adults (see Supplementary Data 9). In some cases, we were unable to sample adult specimens of the same species as the corresponding immature specimen; for these taxa, we selected an adult from the same genus whose skull shape and size were closest to the original species (see Supplementary Data 9).

Specimens collected for this study were obtained from the University of Cambridge Museum of Zoology, the Natural History Museum, Tring, the Natural History Museum of Fribourg, and the National History Museum of Bern. These specimens were scanned using at the Cambridge Biotomography Centre using a Nikon XTEK H 225 ST MicroCT scanner (µCT). Specimens from Fribourg and Bern were scanned using a Bruker Skyscan 2211 at the Geoscience Department of the University of Fribourg, Switzerland Scans of other species were downloaded from the online repository MorphoSource (https://www.morphosource.org/; see Supplementary Data 9 for a detailed list of specimens).

All 3D reconstructions of the PPC bones were obtained from image stacks using the software MIMICS v20 (3D 157 Medical Image Processing Software, Materialize, Leuven, Belgium). Due to extreme bone fusion in some adult specimens, the suture between the rostral portion of the palatine and the surrounding bones (i.e, premaxillary and maxillary) was not always clearly visible. For these specimens, we segmented the rostral portion of the palatine bone based on variation in bone porosity and density, and used immature specimens as a reference as their palate bones remained unfused (see Extended Data Fig. 6).

Because the vomer was reduced or vestigial (n = 2) or even absent (n = 17) in 19 bird species in our sample, we decided not to include this bone in our dataset. Because the palatine and pterygoid are key for the biomechanics of cranial kinesis ^2^, we decided to maximise ecological diversity by sampling as many species as possible, which would not have been the case if the vomer had been included.

During ontogeny, most neognath pterygoids undergo a process of division, named ‘pterygoid segmentation’ ^53^, resulting in the generation of a transient independent hemipterygoid positioned between the palatine and pterygoid in immature specimens. The hemipterygoid then fuses with the caudal end of the palatine in adults^21,53,54,65^. To guarantee comparison between homologous structures, we defined the adult condition, where the hemipterygoid is completely fused to the palatine, as the reference morphology for our analyses. Therefore, for each immature specimen, the hemipterygoid was considered as part of the palatine and included in the palatine landmark scheme (see below for more details). In immature and adult Anseriformes, the anterior region of the pterygoid bears a pronounced rostral process, which has been recognised as homologous to the unsegmented hemipterygoid of immature Neoaves. Therefore, for homology reasons, we have decided to consider this region of the Anseriform pterygoid as part of the palatine, as for the neoavian hemipterygoid.

### Three-dimensional Geometric Morphometrics

As the palatine and pterygoid articulate in intact bird skulls, characterising the shape of the full PPC could be disrupted by bone displacements, especially in osteological preparations of specimens. This is also the case for immature specimens where the bones are not yet in contact due to ongoing ossification. To enable accurate quantification of the full PPC, we followed the approach of Thomas et al., 2023 to generate ‘data blocks’ by landmarking each bone independently and combining these data blocks by scaling the shape configuration for specimen comparisons ^66–68^.

The PPC is a bilaterally symmetrical complex with paired palatines and pterygoids. We were primarily interested in comparisons of the morphology of the palatine with the pterygoid, and as such only focused on one side of the PPC (the left). For species whose left PPC elements were missing or damaged, we mirrored the right bone using Blender (version 3.6).

A set of six anatomical landmarks and eight curves were digitized on the 3D meshes of the palatine, as well as six anatomical landmarks and nine curves on 3D meshes of the pterygoid (see Extended Data Fig. 7 and SI, Supplementary Table 1). All landmarks were manually placed using the software Stratovan Checkpoint, and the curve semi-landmarks situated between landmarks were resampled following previous protocols ^69^ (Divet et al. 2016, see Supplementary Information for relevant code). Sliding (minimising bending energy) and Generalized Procrustes Analysis (GPA),were carried out in the R statistical environment using morphoBlocks ^68^ (Thomas et al., 2021, see Code Availability section).

### Multivariate and phylogenetic analysis

All multivariate and phylogenetic comparative analyses were conducted in the R statistical environment ^70^. For details regarding code, see Code Availability section.

#### Morphological variation of the PPC

To explore morphological variation of the PPC, we conducted a principal components analysis (PCA) (see Extended Data Fig. 1) on each bone independently, and of the complex as a whole.

To estimate morphological disparity in each clade (see above), we calculated Procrustes variance using the *morphol.disparity* function (Geomorph R package^66^). High values of Procrustes variance indicate high morphological disparity according to the whole landmark conformation in the entire sample. To test for differences in PPC morphology between clades, we performed a Procrustes ANOVA using *procD.lm* in the geomorph R package^66^. Then, to assess which clades exhibited significant shape differences, we performed pairwise comparisons using the *pairwise* function from the RRPP R package^71^, where a greater pairwise distance value indicates a greater degree of morphological differentiation between two groups. These tests were conducted on the bony complex as a whole (see Fig. 3) and on each individual bone (see Extended Data Fig. 1).

To evaluate if PPC morphology is correlated with phylogeny, we estimated phylogenetic signal in our Procrustes shape variables, using the *physignal* function in the geomorph R package^66^. Phylogenetic signal was estimated separately on immature adult specimens.

#### Ontogenetic trajectories of the PPC

Morphological variation during ontogeny is strongly associated with size changes^72^. Therefore, most ontogenetic studies to date have focused on the amount of shape variation correlated with size variation (i.e. ontogenetic allometry^28,61,62^). Because our results showed that allometry explained only a small portion of the variation in our data (Supplementary Data 6), we did not undertake further analyses correcting for allometry. To compare ontogenetic trajectories and patterns within and between all major bird clades investigated, we computed three ontogenetic parameters. First, we extracted Euclidean distances between immature and adult specimens of each species as an estimate of the amount of morphological change during ontogeny. Second, because an ontogenetic trajectory can be defined as a vector between the immature and the adult of the same species, we quantified differences in ontogenetic trajectories by comparing vector angles between each species (see Extended Data Fig. 8). We computed angles using the dot product method commonly used to extract this metric between two vectors^73,74^. As a result, the greater the angle between ontogenetic vectors, the greater the difference between ontogenetic trajectories.

Third, as angle measurements do not reveal whether the ontogenetic trajectories between two species are divergent (i.e. shape differences between the adults of different species are more different than shape differences between immatures) or convergent (i.e. the opposite case), we estimated divergence and convergence between ontogenetic trajectories (see Extended Data Fig. 8). For each pair of species, we first calculated the Euclidean distance between immatures (d_1_) and between adults (d_2_), then subtracted the adult-to-immature distance (*d_1-2_*= d_1_-d_2_) (see Extended Data Fig. 8). If *d_1-2_ >0*, the shape differences among the adults are less different than the shape differences among the immatures, indicating a case of convergence; If *d_1-2_ <0*, the shape differences among the adults are more different than those of the immatures, indicating a case of divergence. If *d_1-2_ ∼ 0*, the ontogenetic trajectories can be considered as more or less parallel, indicating a lack of both ontogenetic convergence and divergence.

#### Heterochronic tests

We analyzed evolutionary changes in ontogenetic trajectories following the framework proposed by Alberch et al., 1979^72^, which has since has been widely applied across various phylogenetic groups squamates^27,28,75^, birds^36^, non-avian dinosaurs^76^, or mammals^77^. This approach focuses on examining the relationship between shape and size changes throughout ontogeny. Differences in these relationships across different taxa therefore provide evidence for, and allow us to distinguish between, different heterochronic and non-heterochronic mechanisms shaping ontogenetic differences throughout the evolutionary history of a clade.

Because we lack statistical power to evaluate these relationships between individual species (species-level ontogenies are only one immature and one adult individual) we evaluate these relationships at the clade level ^28^ . This provides insights on signatures of the evolution of ontogenies manifesting at the clade-level. We repeated this set of analyses for the whole PPC complex, as well as the pterygoid and palatine in isolation.

To evaluate which linear models best explained the variation in our morphometric datasets (i.e., Paleognathae vs Neognathae or Paleognathae vs all the Neognathae subgroups), we evaluated the relative fit of different allometric models (see Extended Data Fig. 5) by using the R function ‘model-comparison’ in the RRPP package^78^. We used the -likelihoods and penalty parameters to compare these models, and performed all the downstream tests using the model with the lowest log-likelihood.

To evaluate whether different clades exhibit similar allometries we performed a Homogeneity of Slopes test using the ‘procD.lm’ and ‘anova’ functions in the geomorph^66^ and RRPP packages^78^. When ‘procD.lm’ yielded significant results, indicating differences in allometric slopes, we ran the ‘Het2’ test function^28^ (see detailed function in Ollonen et al., 2024) to test whether the two groups being compared exhibit evidence of neoteny/acceleration (i.e. no difference in slopes) or whether the trajectories are convergent or divergent (i.e. significant differences between slopes).

When the results of ‘procD.lm’ were not significant, we performed a test of homogeneity of intercepts called ‘Het1’^28^ (see detailed function in Ollonen et al., 2024) to test whether the slopes of the two groups under comparison overlap (non-significant p-value) or not (significant p-value). In the event that the slopes do not overlap, it can be concluded that the slopes are parallel. However, if the slopes overlap, we used the peram.test ^28^ (see detailed function in Ollon et al., 2024) to test whether the morphology of the adults between the two groups differ (significant p-value) or not (non-significant p-value). If the results are significant, they suggest the presence of post-displacement, progenesis/hypermorphosis, pre-displacement or neomorphic ontogenetic change depending on the relationship between the two slopes.

#### Phylogenetic context of PPC ontogeny

To test the correlation between developmental mode and ontogenetic parameters (see above), we used a time-calibrated phylogeny from previously published trees^35^. We pruned the original 9,993 species in the original phylogeny^35^ to match our 70-species sample using the R package *ape* ^79^ . To follow the most up-to-date hypothesis of Palaeognathae phylogeny, we modified the tree topology for the group^80^ on our backbone phylogeny.

To compare ontogenetic trajectories within a phylogenetic framework, we needed to estimate a single value per species for the angle and divergence/convergence values. Thus, we first estimated a reference ontogenetic trajectory against which to compare all our species. To do so, we used *gm.prcomp* to reconstruct the ancestral PPC shape for crown group birds (i.e. the last common ancestor of Palaeognathae and Neognathae). To reconstruct ancestral ontogenetic trajectories, we estimated an immature ancestral shape using only immature specimens, and an adult ancestral shape, using only adult specimens. Then, we calculated angles and divergence/convergence values between the hypothetical ancestral ontogeny and descendant ontogenies^36^. These new angles and divergence/convergence values were used to test the relationship between ontogenetic parameters and developmental mode (see below).

To characterise variation in avian developmental patterns, Ducatez and Field (2021)^30^ conducted several PCoA analyses on qualitative traits used to assess developmental mode^29,51^. We used the chick PC scores (ChickPC1 and ChickPC2) results from Ducatez and Field (2021)^30^ as a quantitative estimate of developmental mode along the altricial – precocial spectrum.

To quantify patterns of ontogenetic trajectories, we conducted ordinary least squares (OLS) regressions of ontogenetic parameters and developmental mode using procD.lm in geomorph^66^. We tested whether ontogenetic distances, angles, and divergence/converge values were correlated with developmental mode, then conducted similar regressions within a phylogenetic comparative framework using phylogenetic generalized least-squares (PGLS) using procD.lm in geomorph^66^.

## Acknowledgements

We are indebted to Matt Lowe (UMZC) and Judith White (NHM) for assistance with specimens, and Keturah Smithson (Cambridge Biotomography Centre) for CT scanning assistance. We would like to thank Michel Beaud and Boris Baeriswyl (Musée d’Histoire Naturelle Fribourg), Manuel Schweizer and Reto Hagmann (Naturhistorisches Museum Bern) for giving access to collection specimens and Sharon Grant for giving access to MorphoSource specimens. We thank Annabel Hunt (Cambridge) for insight into hemipterygoid homology guiding our landmark applications. This work was funded by UKRI grant MR/X015130/1 and the SNSF grant P500PN_214284. For the purpose of open access, the authors have applied a Creative Commons Attribution (CC BY) licence to any Author Accepted Manuscript version arising.

## Extended Data Figure legends (up to 10)

**Extended Data Figure 1.**
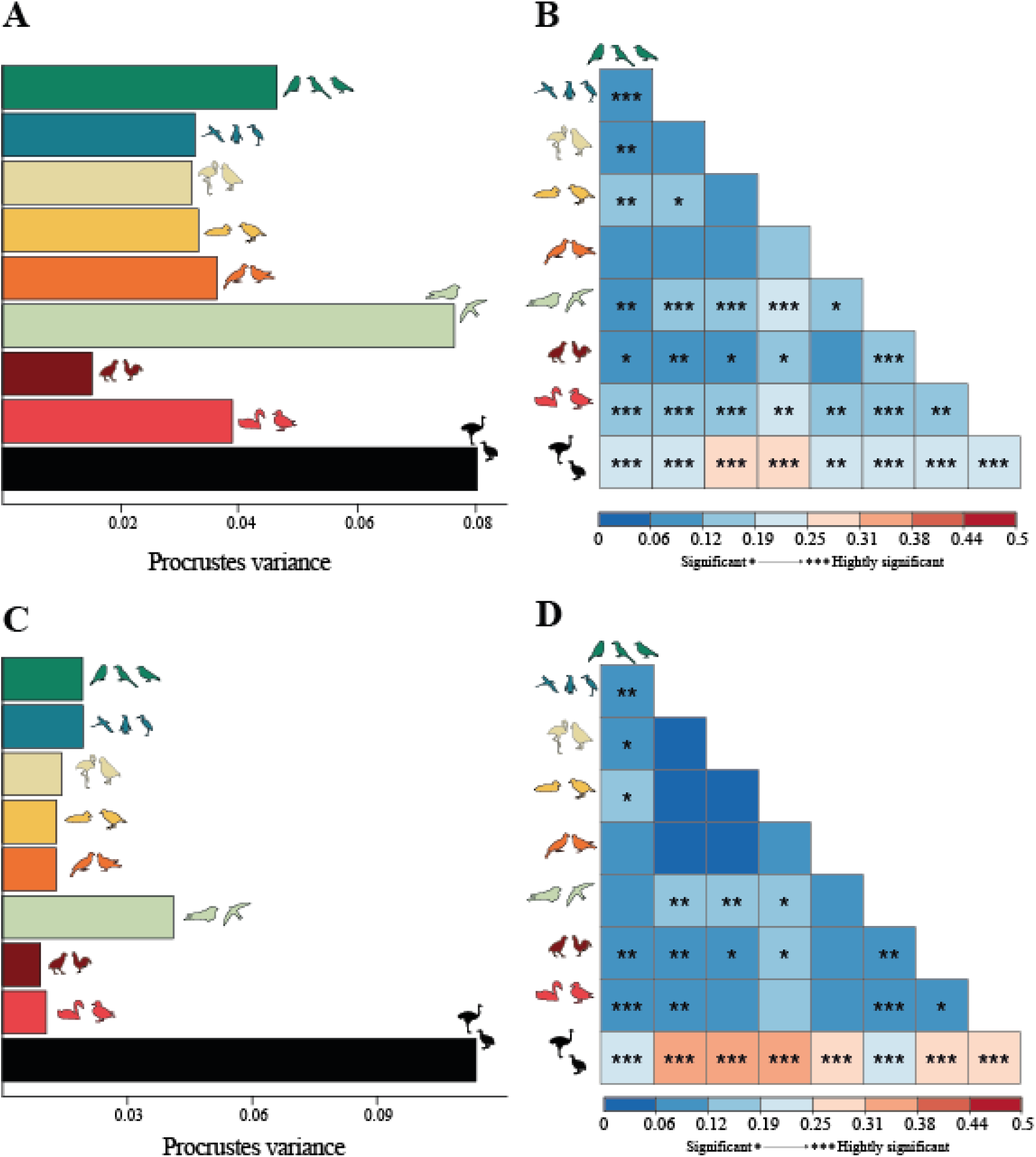
Morphological disparity and pairwise group comparisons by bone. Data for palatine in (A) and (B); data for pterygoid in (C) and (D). Bar plot showing Procrustes variance per avian subclade investigated (A) and correlation plot pairwise distances between each subgroup, with asterisks indicating significance (* = 0.05>P>0.01; ** =0.01>P>0.001; *** 0.001>P) (B). Same plots are shown for the pterygoid (C and D). Subclade coloursfollow Figure 1. See Supplementary Data 3 for detailed results.

**Extended Data Figure 2.**
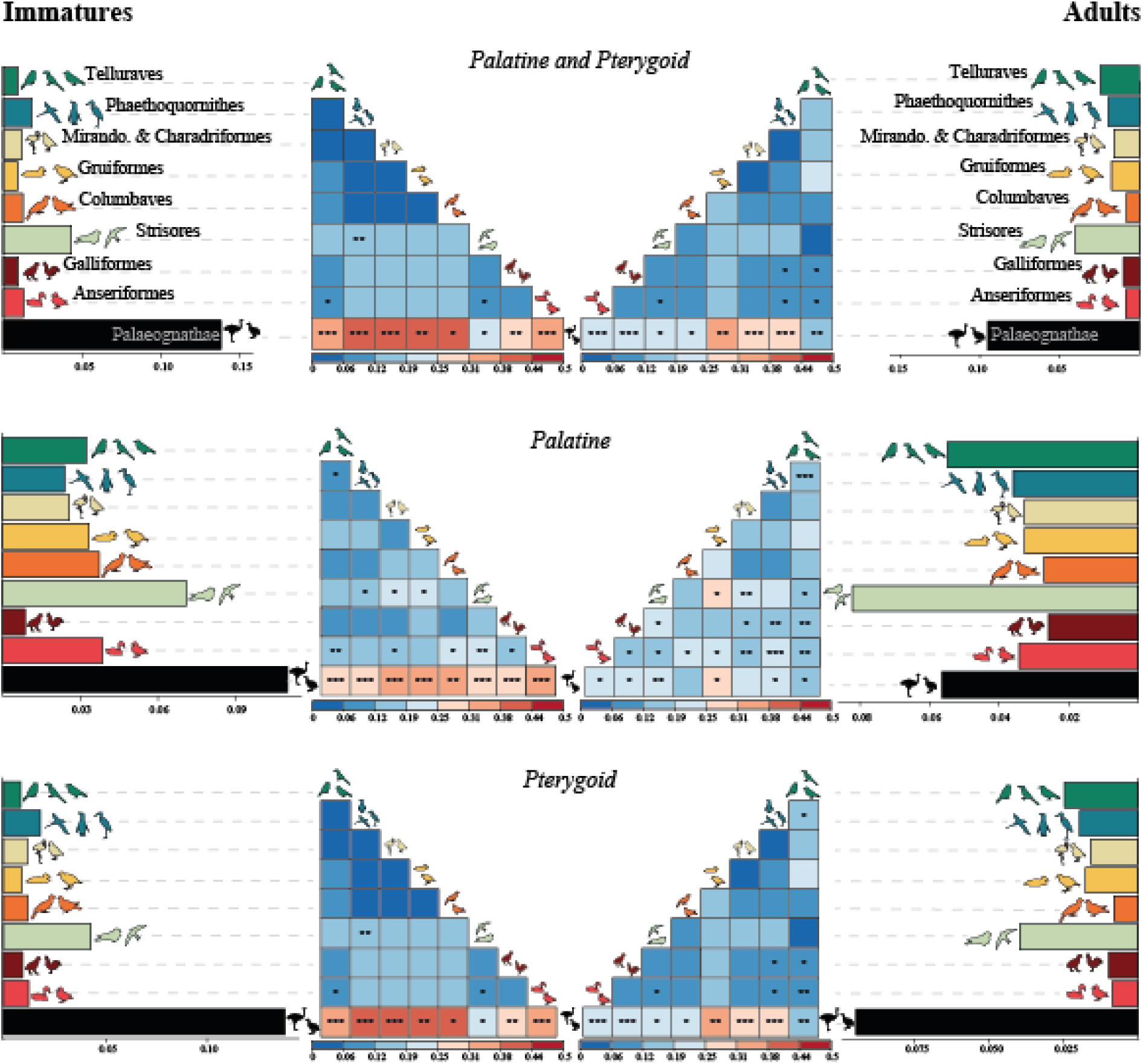
Morphological disparity and pairwise group comparisons by bone and stage. Bar plot showing Procrustes variance per avian subclade investigated (left and right columns represent immatures and adults, respectively) and correlation plot of the pairwise distances between each subgroup, with asterisks indicating significance (* = 0.05>P>0.01; ** =0.01>P>0.001; *** 0.001>P). The two central columns represent immatures (left) and adults (right) for the full PPC complex (top), the palatine only (middle) and the pterygoid only (bottom). Subclade colours follow Figure 1. See Supplementary Data 1 for detailed results tables.

**Extended Data Figure 3.**
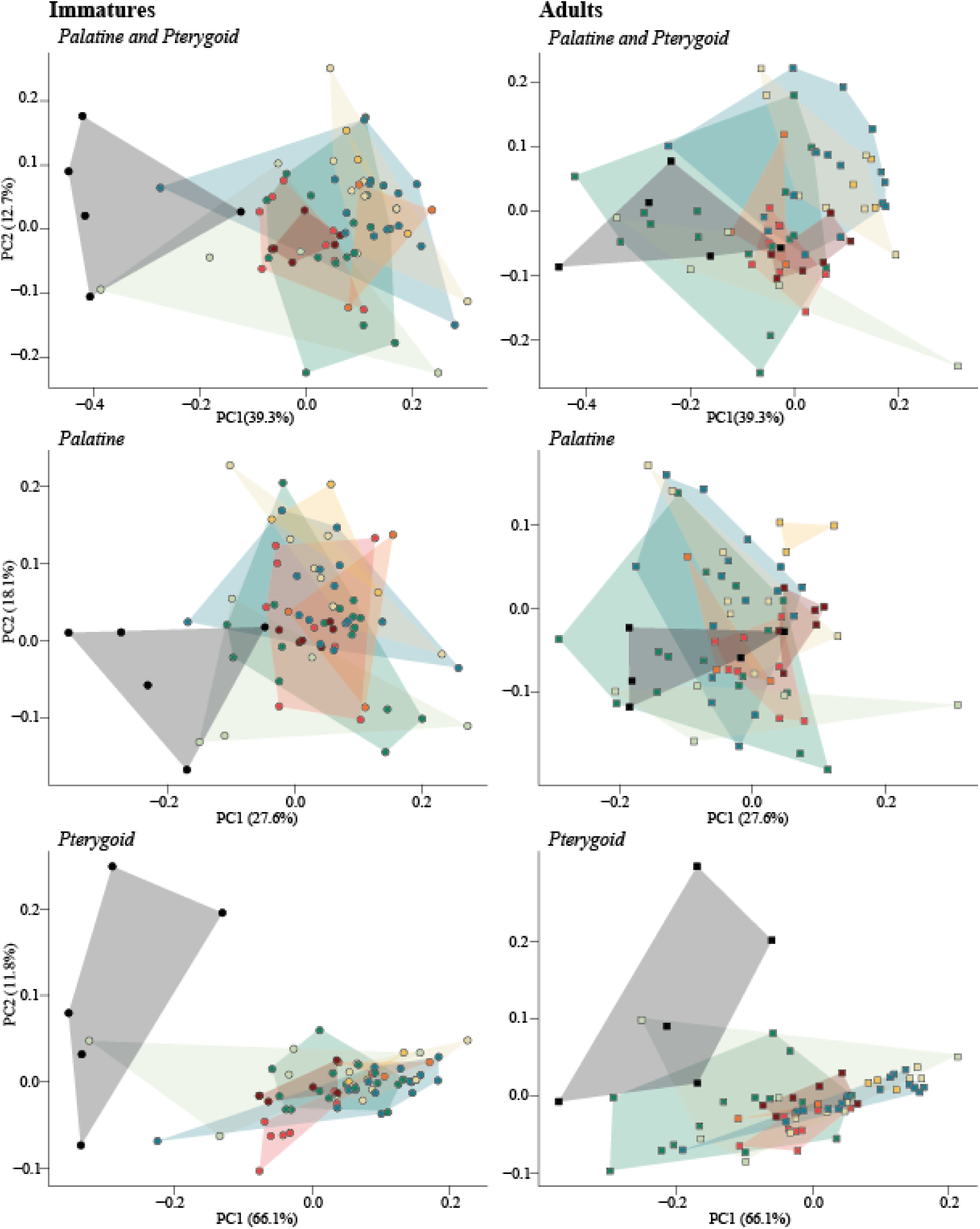
Principal Component Analysis (PCA) plots by bone and stage. Graphs show PCA morphospace based on the first two PCs of the full complex (as in Figure 3A), the palatine alone (as in Figure 4A) and the pterygoid alone (as in Figure 4B). In each case, PCAs for immatures are shown on the left and adults on the right. Circles indicate immature specimens and squares indicate adults. Subclade colours follow Figure 1. See Supplementary Data 10 for detailed results tables.

**Extended Data Figure 4.**
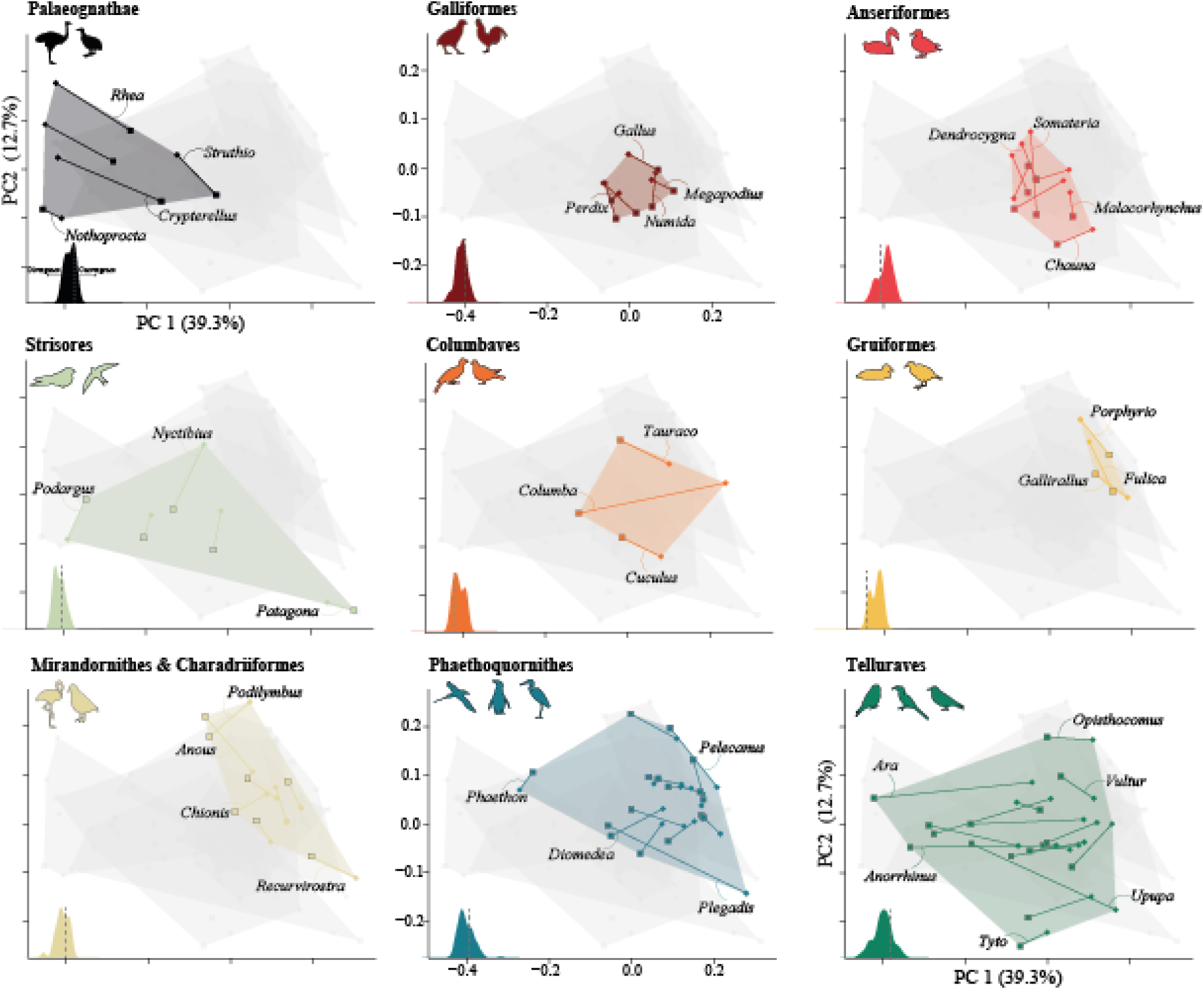
Principal Component Analysis (PCA) per avian subclade investigated. Graphs show PCA morphospaces based on the first two PCs, as in Figure 3. Circles indicate immature specimens and squares indicate adult specimens, each line links immatures and adults of the same species. Inset plot on the bottom left of each PC plot corresponds to the frequency plot in Figure 5.

**Extended Data Figure 5.**
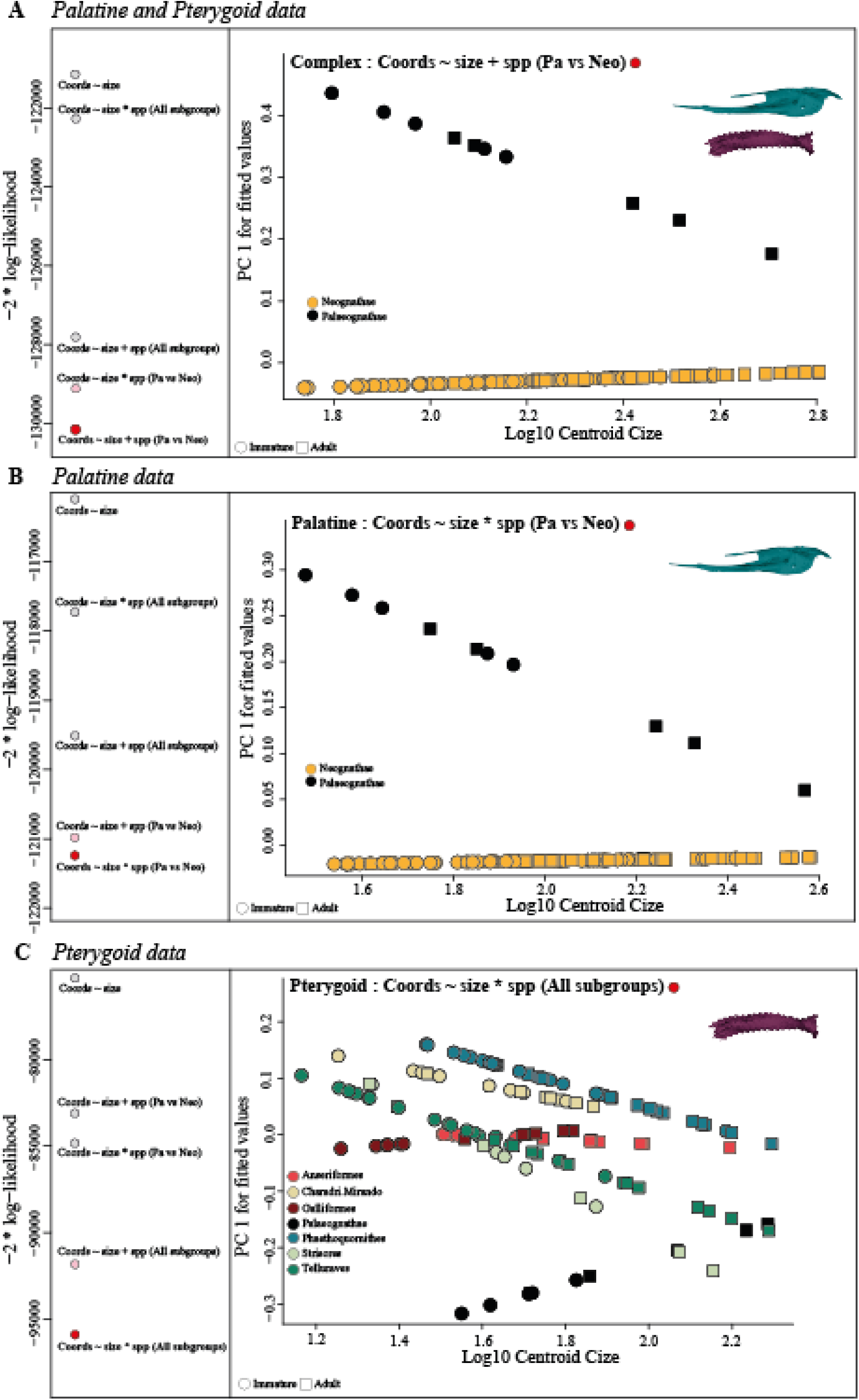
Heterochronic tests within the PPC for A) palatine alone B) pterygoid alone, and C) the full PPC complex (combined palatine and pterygoid). For each plot the left-hand column represents the graphical results of model comparisons to test which models are most appropriate for our dataset. Models compared are: correlation between shape and size (coords ∼ size); correlation between shape and the interaction between size and phylogenetic groups (coords ∼ size*spp); correlation between shape and size and phylogenetic groups as independent variables (coords ∼ size + spp). Each model is run to compare Palaeognathae and Neognathae (Pa vs Neo) as well as Palaeognathae with the major Neognathae subgroups investigated. The best model (per AIC) is represented by the lowest log-likelihood value, indicated by a red circle. Grey circles indicate the weakest correlation, and pink circles show the second-best correlation tested. For each graph, the right-hand column represents the bivariate plot between fitted values and size for the best model comparison.

**Extended Data Figure 6.**
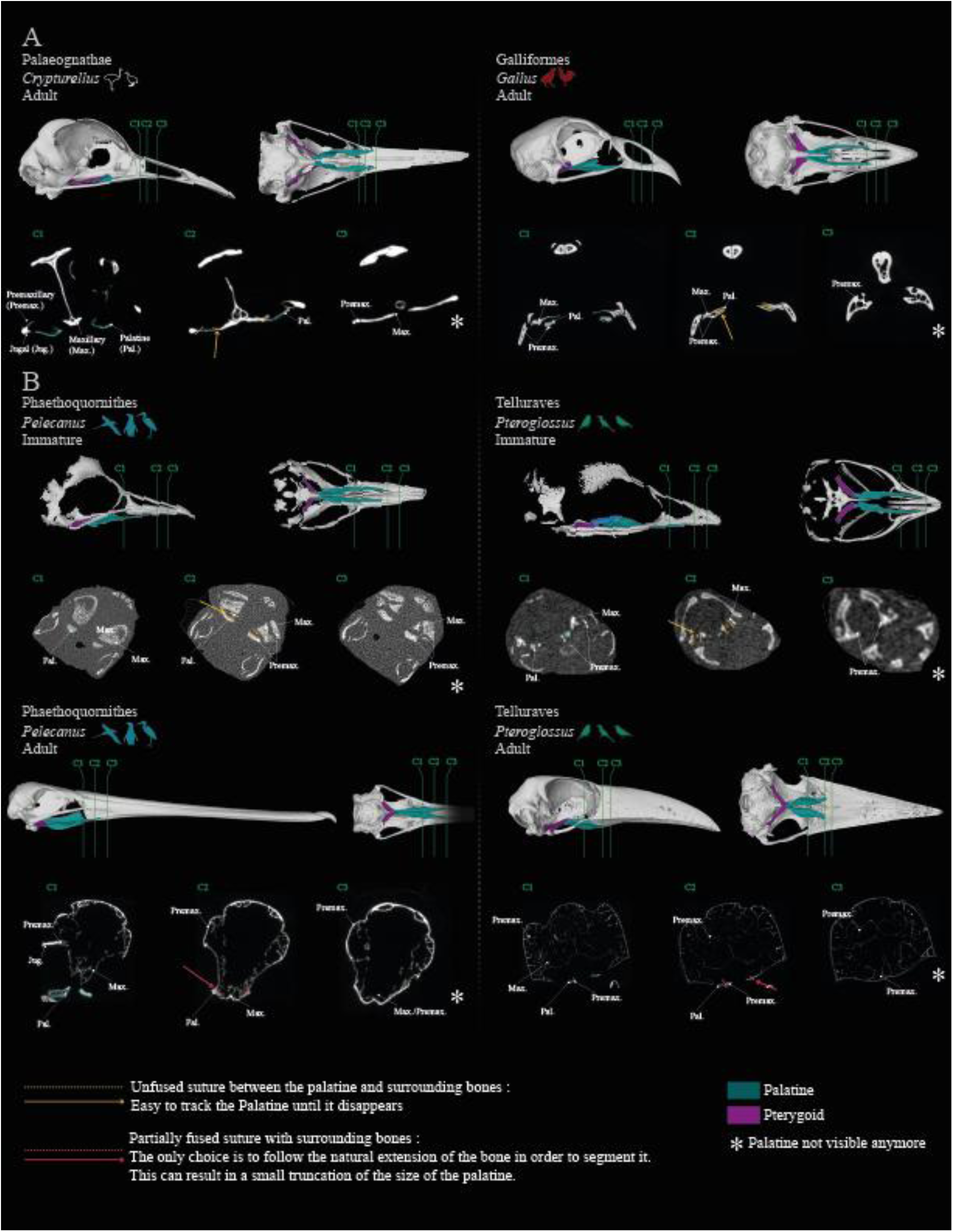
Anatomical plate illustrating the digital segmentation of the rostral-most portion of the palatine. Lateral and ventral cranium views of (A) adult *Crypturellus tataupa* (left) and *Gallus gallus* (right) and (B) immature *Pelecanus philippensis* and adult *Pelecanus occidentalis* (left) and immature and adult *Pteroglossus viridis* (right). For each specimen, three sections (C1, C2, C3) are shown in coronal view to highlight the presence of specific bones (premaxilla, maxilla, jugal and palatine). Subclade colours follow Figure 1.

**Extended Data Figure 7.**
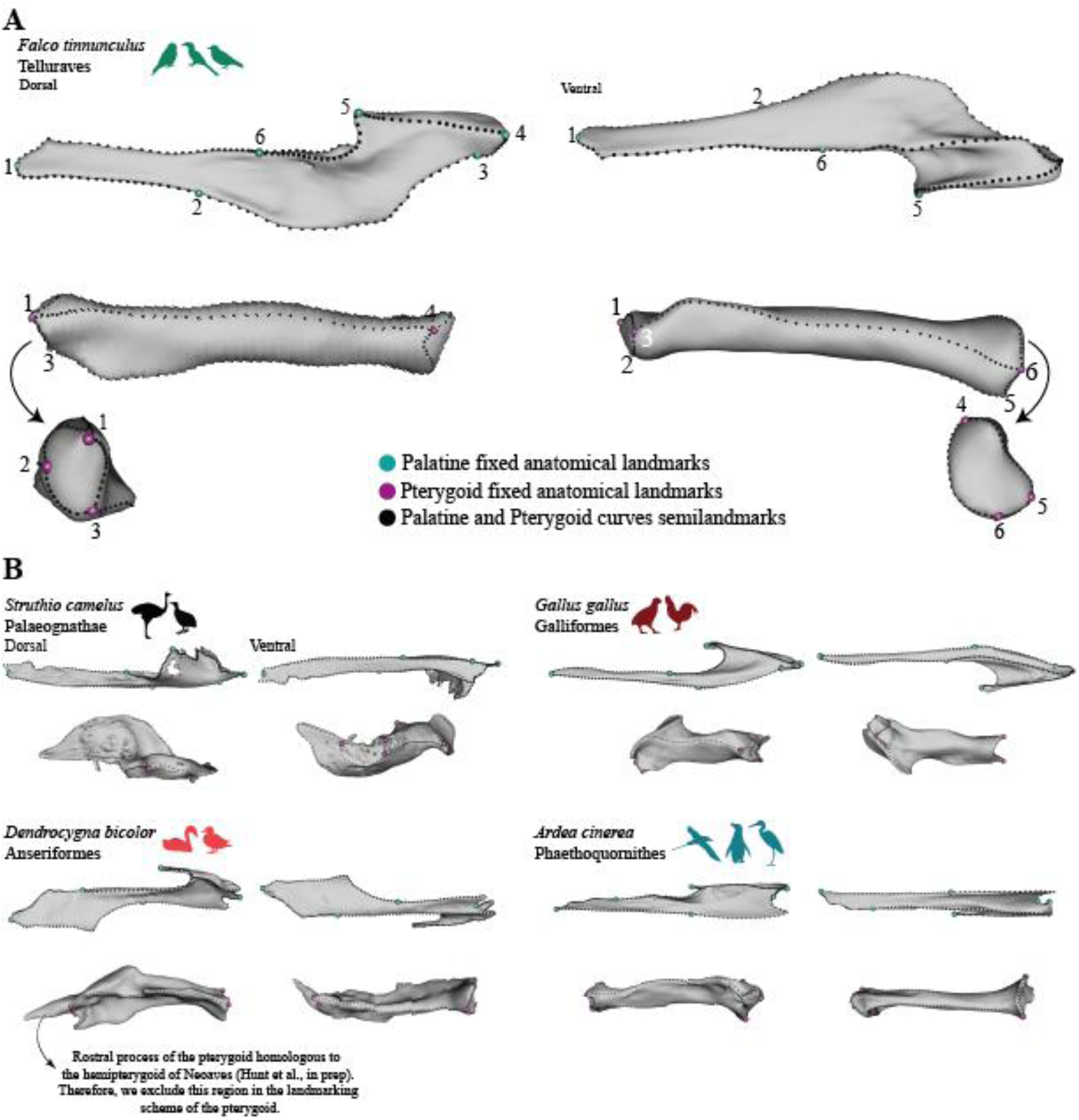
Anatomical plate illustrating landmark placement used in this study. A) Detailed position of the palatine and pterygoid landmarks. B) Overview of the placement of the palatine and pterygoid landmarks in species belonging to various groups. See Supplementary Table 1 for detailed landmark positions.

**Extended Data Figure 8.**
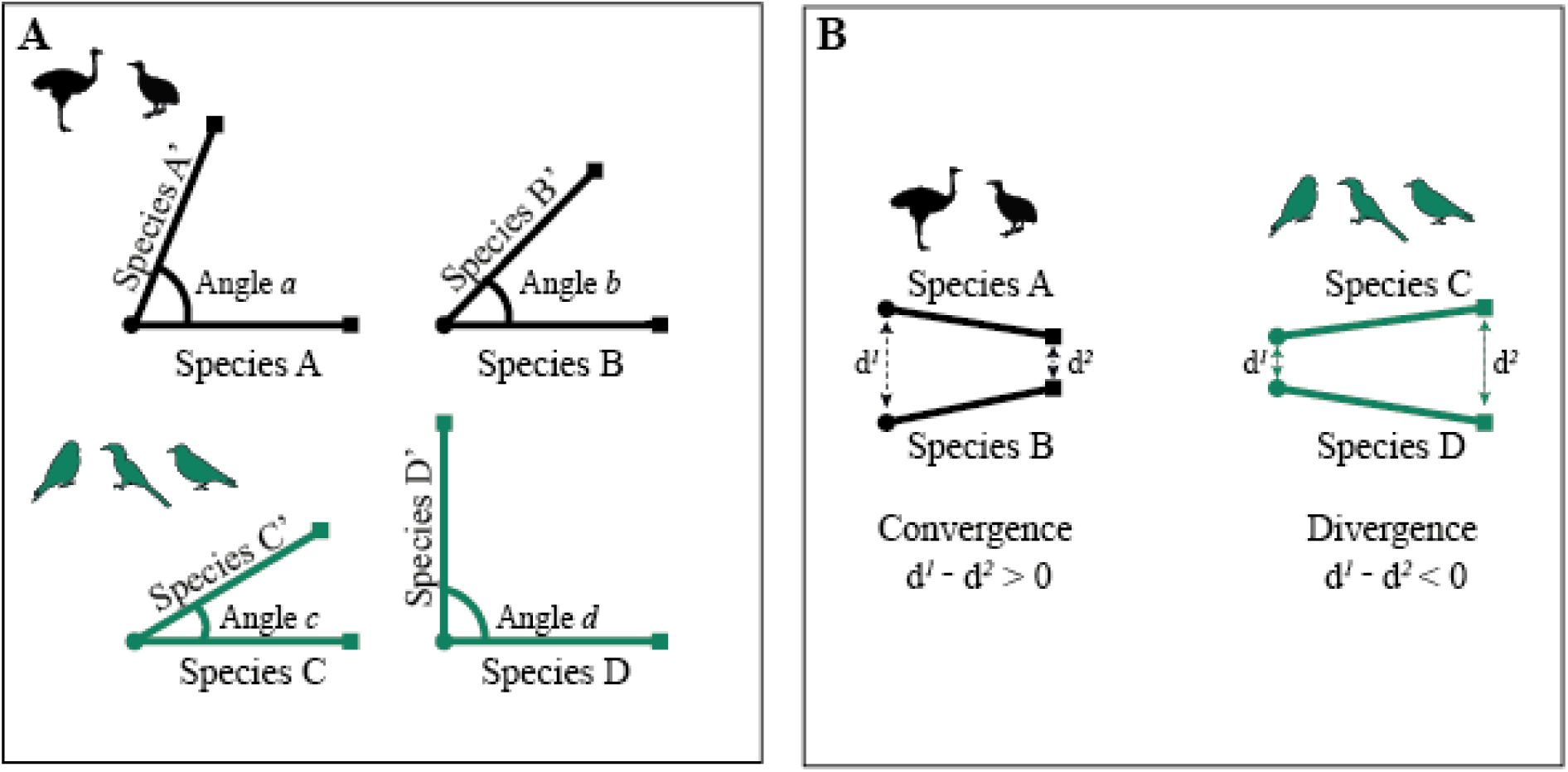
Graphical representation of the method used to compare ontogenetic trajectories. A) Hypothetical angle of comparison of ontogenetic trajectories between species of Palaeognathae (black) and Telluraves (green). Circles indicate immature specimens and squares adult specimens; lines indicate different species. Different angles represent hypothetical variation in angles between species. B) Hypothetical convergence and divergence in comparisons between species of Palaeognathae and Telluraves. d1 represents the distance between immature specimens and d2 the distance between adult specimens of two different species.

