## Supplementary information for "Developmental underpinnings of morphological disparity in the avian bony palate"

### Supplementary Tables

| | |
|---|---|
| **Palatine landmark scheme** | |
| **Anatomical landmarks** | |
| **1** | Most anterior point of the maxillary process of the palatine |
| **2** | Most anterior-lateral point of the pars lateralis |
| **3** | Most posterior - lateral point of the pars lateralis |
| **4** | Most posterior point of the processus pterygoideus of the palatine (dorsally) |
| **5** | Most anterior point of the processus choanalis |
| **6** | Most anterior part of the ridge of the pars choanalis |
| **Curves landmarks** | |
| **C1** | Laeral marging of the maxillary process (Between 1 and 2) |
| **C2** | Medial margin of the maxillay process (Between 1 and 6) |
| **C3** | Lateral margin of the pars lateralis (Between 2 and 3) |
| **C4** | Medial margin of the pars lateralis (Between 6 and 4) |
| **C5** | Lateral margin of the porcessus pterygoideus (Between 3 and 4) |
| **C6** | Medio-dorsal margin of the processus choanalis (Between 4 and 5) |
| **C7** | Anterior curvature of the processus choanalis (Between 5 and 6) |
| **C8** | Medio-ventral margin of the processus choanalis (Between 4 and 5) |
| **Pterygoid landmark scheme** | |
| **Anatomical landmarks** | |
| **1** | Most anterior point of the pterygoid feet that join the most dorsal crest |
| **2** | Most anterior point of the pterygoid feet that join the most medial / mediolateral crest |
| **3** | Most antero-ventral point of the pterygoid feet that join the most ventral / ventrolateral crest |
| **4** | Most postero-medial point of the quadrate process that join the dorsal crest |
| **5** | Most postero-medial point of the quadrate process that join the medial crest |
| **6** | Most posterior-ventral point of the quadrate process that join the ventral crest |
| **Curves landmarks** | |
| **C1** | Shape of the pterygoid feet (Between 1 and 2) |
| **C2** | Shape of the pterygoid feet (Between 2 and 3) |
| **C3** | Shape of the pterygoid feet (Between 3 and 1) |
| **C4** | Shape of the quadrate process (Between 4 and 5) |
| **C5** | Shape of the quadrate process (Between 5 and 6) |
| **C6** | Shape of the quadrate process (Between 6 and 4) |
| **C7** | Left crest (Between 1 and 4) |
| **C8** | Right crest (Between 2 and 5) |
| **C9** | Ventral crest (Between 3 and 6) |

**Supplementary Table 1: Detailed description of the landmark scheme for the palatine and the pterygoid**

### Supplementary Results

#### Additional results – Quantifying morphological disparity among major bird clades

Details about neognath subclades: All neognath subclades exhibit substantial overlap in along PC1-PC2 morphospace, with Galloanserae clustering near the centre. While most Phaethoquornithes (waterbirds) occupy the positive side of PC2, the great majority of Telluraves plot along the negative side of PC2. Furthermore, while Mirandornithes (flamingos and grebes), Charadriiformes (shorebirds) and Gruiformes (cranes and allies) generally overlap with Phaethoquornithes, Strisores mainly overlap with Telluraves (Figs. 3A-B).

#### Additional results – Ontogenetic trajectories across avian phylogeny

- ***Group-specific shape differences in ontogenetic trajectories***

Details about specific subclades angles: Galliformes and Strisores exhibit only limited intraclade variation (with variation mainly clustering around 90°), whereas most other neognath subclades exhibit a wider range of values (e.g., Fig. 5A; Supplementary Data 4). Although represented by only a sparse taxon sample, the neognath subclade Gruiformes exhibits bimodal groupings clustered around 54° and 105°, while all investigated representatives of the major subclade Columbaves cluster around 55° (Fig. 5A, Supplementary Data 4).

- ***Ontogenetic shape divergence and convergence among birds***

Details about Galloanserae patterns: However, in Anseriformes, we detect the opposite pattern, in which PPC shape tends to converge among species during ontogeny (Fig. 4C) (i.e. adults of different species are more geometrically similar than immatures are). Although Galliformes show more ontogenetic divergence than Anseriformes, comparison of ontogenetic trajectories within Galloanserae (Galliformes + Anseriformes) shows a peak value at 0, indicating a tendency towards parallel trajectories. We also detect a convergent pattern in Columbaves and Gruiformes; however, these clades are sparsely sampled in our dataset and may not reflect broader patterns within these taxa.

- **Non-heterochronic ontogenetic variation in the avian PPC**

Details about models tested: Based on model comparisons (see Materials and Methods section for details), we found that comparisons of ontogenetic trajectories between Paleognathae and Neognathae were the most effective basis for testing the hypothesis of heterochrony for the full

PPC complex as well as the palatine in isolation (Extended Data Fig.5, Supplementary Data 6). However, for the pterygoid, the best model for testing heterochrony was to compare ontogenetic allometries between Palaeognathae and the major subgroups of Neognathae (Extended Data Fig.5C, Supplementary Data 6).

Details about specific pattens in neognathae: Within Neognathae, Anseriformes and Galliformes exhibit significant differences in their ontogenetic trajectories with respect to select other neognath subclades (Anseriformes vs Strisores and Telluraves; Galliformes vs Phaethoquirnithes, Strisores and Telluraves). Anseriform and galliform trajectories do not overlap in shape-space (Thf2: significant results, Extended Data Fig.5C), indicating that their ontogenetic trajectories converge towards each other during post-hatching ontogeny (Extended Data Fig.5C). No other neognath groups exhibit significant differences in their ontogenetic trajectories. While the ontogenetic trajectories of most neognath subclades do not overlap in size-shape (Thf1: significant), indicating parallel slope, those of Mirandornithes-Charadriiformes vs Phaethoquornithes and Strisores vs Telluraves overlap in size-shape (Thf1: non-significant). However, we did not detect any ontogenetic scaling (Peram test non-significant) indicating that, although their slopes overlap, they are best considered parallel and of the same length.

### Supplementary Discussion

#### Evolutionary patterns of palatal ontogeny across living birds

Our results reveal a significant degree of neomorphic ontogenetic variation across main lineages but also within specific groups. For instance, within Palaeognathae, the two sampled Tinamiformes, *Nothoprocta* and *Crypturellus,* present the most unique and divergent ontogenetic trajectories (Supplementary data). Their elongated pterygoids are very similar yet vary in proportions between immatures and adults, while their palatines, similar in size, are wildly different, and each resembles that of a more distantly related palaeognath instead (ostriches and emus, respectively). Galloanserae, except for a few extreme values (i.e, values of divergence or convergence), exhibit a predominance of parallel ontogenetic trajectories (Fig. 5 B), indicating that both Anseriformes and Galliformes display similar ontogenetic variation. This could be the result of specific phylogenetic and developmental constraints, perhaps related to the absence of pterygoid segmentation in Galloanserae[1].

Neoaves, as a much more diverse clade, shows a greater distance between ontogenetic trajectories (Figure 5.A) leading to higher divergences between lineages. Telluraves, which encompasses most of neoavian diversity, shows the greatest distance between ontogenetic trajectories and the highest values for ontogenetic trajectory direction (Figure 5.A), while other neoavian clades show more idiosyncratic pattern within groups, i.e. each group presents a distinctive pattern straddling the range between Galloanserae and Telluraves.

Within Telluraves,strong morphological convergence—reflected by a large angular differences and higher convergence values—can occur, as demonstrated by *Falco tinnunculus* and *Falco naumanni* (see Supplementary Data 4 and 5). Also, the amount of shape changes is slightly lower between immature and the adult of *F. tinnunculus* than *naumanni*, this results also highlights the potential for substantial ontogenetic variation even among closely related taxa. Notably, our Telluraves sample is predominantly composed of altricial and super-altricial species, whereas *Falco* represents a semi-altricial lineage. As discussed, semi-altricial and semi-precocial birds exhibit less clearly defined ontogenetic patterns compared to the extremes of the spectrum (i.e., super-altricial to super-precocial). Investigating the detailed ontogenetic development of other semi-altricial Telluraves would be valuable for better understanding the developmental implications of semi-altriciality within this clade. Morever, to fully elucidate the relationship between broad evolutionary trends and lineage-specific developmental pathways, future research should include more comprehensive taxonomic sampling and in-depth ontogenetic analyses of closely related species.
